# Nephron-associated Support Cell Transcriptional Plasticity Expands in Hypertension

**DOI:** 10.64898/2026.01.16.699969

**Authors:** Justin G. McDermott, Bethany L. Goodlett, Shobana Navaneethabalakrishnan, Joseph M. Rutkowski, Brett M. Mitchell

**Affiliations:** Department of Medical Physiology, Texas A&M University Vashisht College of Medicine, Bryan, TX 77807

**Keywords:** stem cell, kidney, hypertension, inflammation, gene expression

## Abstract

Hypertension (HTN) affects over one billion people worldwide and can lead to debilitating cardiovascular and renal conditions if left untreated. Cell death in the kidneys and the inflammation that follows are among the primary effects of chronically elevated blood pressure. There are several cell types throughout the body with immunomodulatory, anti-inflammatory, and pro-regenerative properties that support tissue homeostasis and recent studies have highlighted their therapeutic potential in HTN and kidney-related conditions. In our previous paper, we found a pool of multipotent nephron-associated support cells (SCs) in single-cell RNA sequencing samples of CD31+ and podoplanin+ cells taken from the kidneys of hypertensive mice generated through two mouse models of HTN. Despite remaining roughly constant in number between HTN and control groups, these SCs had 299 differentially expressed genes (p<0.01), 51 and 86 enriched pathways (p<0.01) in the M2 and M5 Molecular Signatures Database gene sets, respectively, and 180 HTN-specific regulons. We also compared lymphatic endothelial cells (LECs) and SCs from HTN and control groups and identified 3636 differentially expressed genes (p<0.01), 537 M2 and 415 M5 enriched pathways (p<0.01), and 218 LEC-specific and 227 SC-specific regulons in the HTN samples. SCs from mice with HTN were more resistant to inflammation-induced changes compared to LECs, and had downregulated stem cell suppressive genes and upregulated genes related to stem cell proliferation and regeneration.

**Graphical Abstract:** Created with BioRender.com

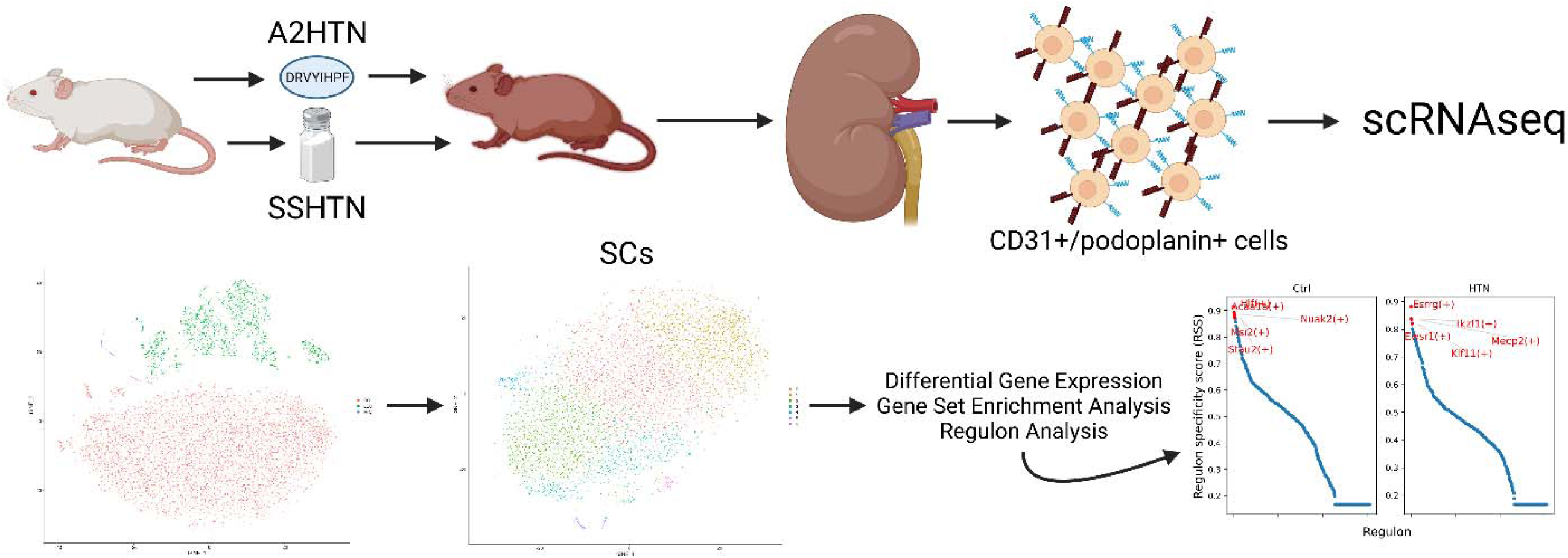

## Introduction

Hypertension (HTN) is a pro-inflammatory condition that affects over 1.2 billion adults worldwide and is the primary risk factor for deadly cardiovascular diseases.^1^ Prolonged HTN causes organ damage and dysfunction of the brain, eyes, blood vessels, reproductive organs, and the kidneys among others. First-line treatments include lifestyle changes and/or medications such as thiazide diuretics, calcium channel blockers, angiotensin converting enzyme inhibitors, and angiotensin receptor blockers according to the JNC 8 guidelines.^2^ However, almost half of patients have uncontrolled HTN mostly due to medication non-compliance, negative side effects of medications, or decreased drug efficacy thus prompting additional medications to be combined.

Kidney damage is a common element of severe HTN that results from excessive blood pressure (BP) impairing the cells that constitute the pressure-sensitive structures throughout the kidneys. A previous review article discusses the pathophysiology of HTN-induced renal damage and therapeutic strategies, including damage to microvasculature in and around nephrons and how standard treatments for HTN only partially address these issues.^3^ This damage leads to an increase in local inflammation, which then contributes to immune cell infiltration and activation and even further damage and inflammation.^4^ Over time, this pro-inflammatory loop can contribute to fibrosis and a decline in kidney function.^5–9^ Many studies have been conducted to better understand how this loop begins and propagates in the context of HTN with the intention of breaking the cycle of damage.

Recently, a handful of studies have evaluated the role of various forms of stem cells, which are generally considered to be anti-inflammatory and can promote healing and regeneration, in HTN and other kidney-focused conditions. Studies involving the injection of adipose or bone marrow derived stem cells (or the extracellular vesicles and factors they release) into hypertensive animals have reported improvements in BP and renal inflammation, damage, and function.^10–13^ Similar findings have also been reported in chronic kidney disease and polycystic kidney disease models, in human kidney transplant patients, and through single nucleus RNA sequencing of mouse kidneys following acute kidney injury.^14–17^

In our previous paper, we performed single cell RNA sequencing (scRNAseq) on CD31+/podoplanin+ renal cells from hypertensive and normotensive mice to better understand the changes that occur in lymphatic endothelial cells (LECs) in response to HTN using the angiotensin II-induced (A2HTN) and salt sensitive (SSHTN) mouse models of HTN.^18^ In addition to LECs, we identified a substantial pool of an unexpected multipotent cell type that underwent a variety of notable changes in hypertensive animals, so we chose to analyze them to better understand the role of these cells, which we dubbed support cells (SCs), as well as the changes they undergo in HTN and their relationship with LECs.

## Methods

### Mouse Models of HTN and Kidney Dissociation

All procedures performed in mice were approved by the Texas A&M University Institutional Animal Care and Use Committee (#2022-0083) and performed at the Texas A&M College of Medicine in accordance with the NIH Guide for the Care and Use of Laboratory Animals. ARRIVE guidelines were followed for conducting this animal study. As described previously, groups of male C57/BL6 mice (n=6 per group to provide sufficient cells and account for natural variation) from Jackson Laboratories were subjected to two common models of HTN (A2HTN and SSHTN), and their respective controls, when the animals were between 10-14 weeks of age following at least one month of acclimatisation.^18–20^ Mice in the A2HTN and A2 control groups were implanted under anesthesia with subcutaneous osmotic pumps, which delivered 3 weeks of either 1,000 ng/kg/min of angiotensin II in saline solution or pure saline. Mice in the SSHTN and SS control groups both received 2 weeks of 0.5 g/L of LNAME by drinking water followed by a washout period of 2 weeks, then a diet of either 4% high salt chow or regular chow for 3 weeks. Following the 3 weeks of either angiotensin II, high salt chow, or their controls, the mice were euthanized with isoflurane and cervical dislocation and the kidneys were removed, then minced and subjected to enzymatic digestion (Collagenase D (2.5 mg/mL, Roche) and Dispase II (1 mg/mL, Sigma-Aldrich)).^19,21^

### Enrichment for CD31+/Podoplanin+ Cells, Library Preparation, and Sequencing

The suspension of kidney cells was filtered with 70 μm and 40 μm filters, then washed twice prior to incubation with biotinylated rat anti-mouse CD31 antibodies (BD Biosciences, catalog number: 553371) and separation with the Invitrogen CELLection Biotin Binder Kit (Thermo Fisher Scientific, catalog number: 11533D), which was repeated with biotinylated goat anti-mouse podoplanin antibodies (R&D Systems, catalog number: BAF3244) in place of the anti-mouse CD31 antibodies. Cells were filtered and washed again, followed by library preparation using the 10X Genomics Chromium Next GEM Single Cell 3′ Reagent Kit V3.1 with its standard protocol. The Agilent High Sensitivity D5000 ScreenTape assay was used to verify each sample’s cDNA, then samples were sequenced using the Illumina NovaSeq 6000 with an S4 flow cell.

### Transcriptome Alignment, Quality Control, Clustering and Identification, and Differential Gene Expression

Cell Ranger (v7.0.0) with the 10mm mouse genome assembly was used for alignment using default parameters. Afterwards, Seurat (v4.1) was used for quality control, clustering of cells, PCA, gene expression plotting, and differential gene expression.^22^ Cells with more than 3,000 genes or fewer than 300 genes were removed along with cells with a mitochondrial gene expression ratio above 10%.^23^ Doublet-Finder (v2.0.3) was also utilized to find and remove potential doublets from the samples.^24^ Based on cell enrichment method and the similarities and differences in gene expression relative to previous studies, clusters were designated as lymphatic endothelial cells, myeloid immune cells, and a multipotent cell type we referred to as support cells (SCs).^25–27^ Differential gene expression was analyzed between cell types and conditions using Seurat’s default parameters with controls from both HTN models being combined into a single control group.

### Gene Set Enrichment and Regulon Analyses

Gene set enrichment analysis (GSEA) using the curated and ontology murine gene sets from the Molecular Signatures Database utilized a combination of FGSEA (v1.20) and msigdbr (v7.5.1) using default parameters.^28^ Regulon analysis was performed using SCENIC (pySCENIC v0.11.2) with default parameters and murine transcription factor motifs to identify positive regulons with differential activity specific to a cell type or condition.^29^

## Results

### Cell Type Identification of SCs

Previously, we identified populations of three distinct cell types from scRNAseq of CD31+/podoplanin+ cells isolated from the kidneys of hypertensive and normotensive mice and analyzed the changes to gene expression in LECs in response to HTN. Using three established murine kidney datasets from NCBI GEO (Janosevic: GSE151658, Conway: GSE140023, Miao: GSE157079), we first attempted to compare our samples and determine the identity of the SCs by mapping them together with established kidney cell types.^30–32^ There was no overlap between the SC population and the three datasets, though there was some minimal overlap with LECs and myeloid immune cells to the dataset produced by Miao et al. (Figure S1).^30^ In order to better understand the identity and role of SCs, we utilized a list of previously established marker genes from other scRNAseq studies of kidney cell types to create gene feature plots (Figures S1C & S2).^30,33–36^

There was significant overlap between markers for podocytes (*Nphs1* and *Nphs2*), proximal tubule (*Slc34a1* and *Slc27a2*), distal tubule (*Slc12a3*), loop of Henle (*Slc8a1* and *Slc12a1*), mesangial (*Prcka* and *Nt5e*), and endothelial (*Pecam1* and *Egfl7*) cell types. Considering the clear distinctions between SCs and cells from the established datasets, the combination of cell type markers that SCs express, and the variety of responses they showed in HTN samples during initial analysis, we designated this population as multipotent nephron-associated support cells. SCs from each group were isolated, reclustered, and then reanalyzed for cluster marker genes (Figures 1 and S4; Tables 1-3 and “Cluster Genes” file Tables S1-5). There was little difference in terms of the spread in TNSE plots of SCs from HTN and control (Ctrl) samples and the number of clusters remained constant from the merged Ctrl group to the merged HTN group.

**Figure 1.**
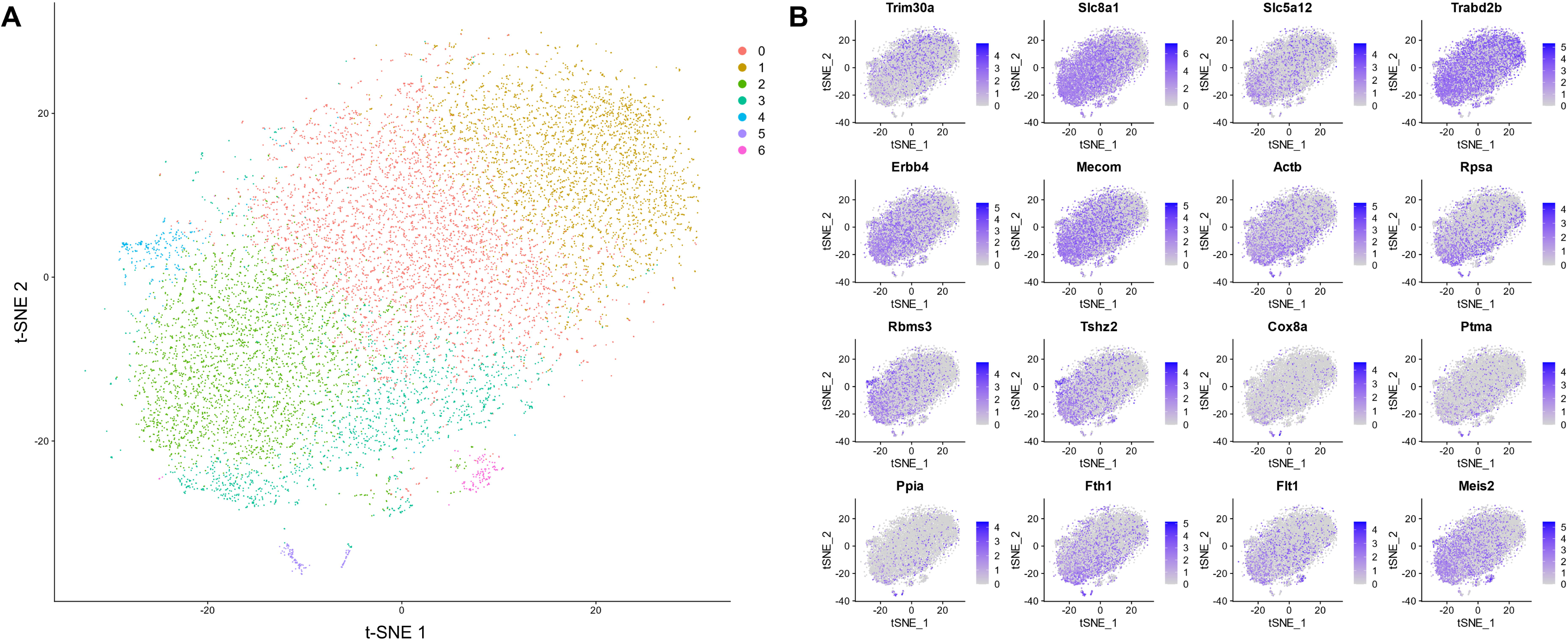
TSNE cluster plot and feature plot of SCs from all samples. The features shown here were the most consistent and selective SC cluster markers across all samples.

**Table 1.**
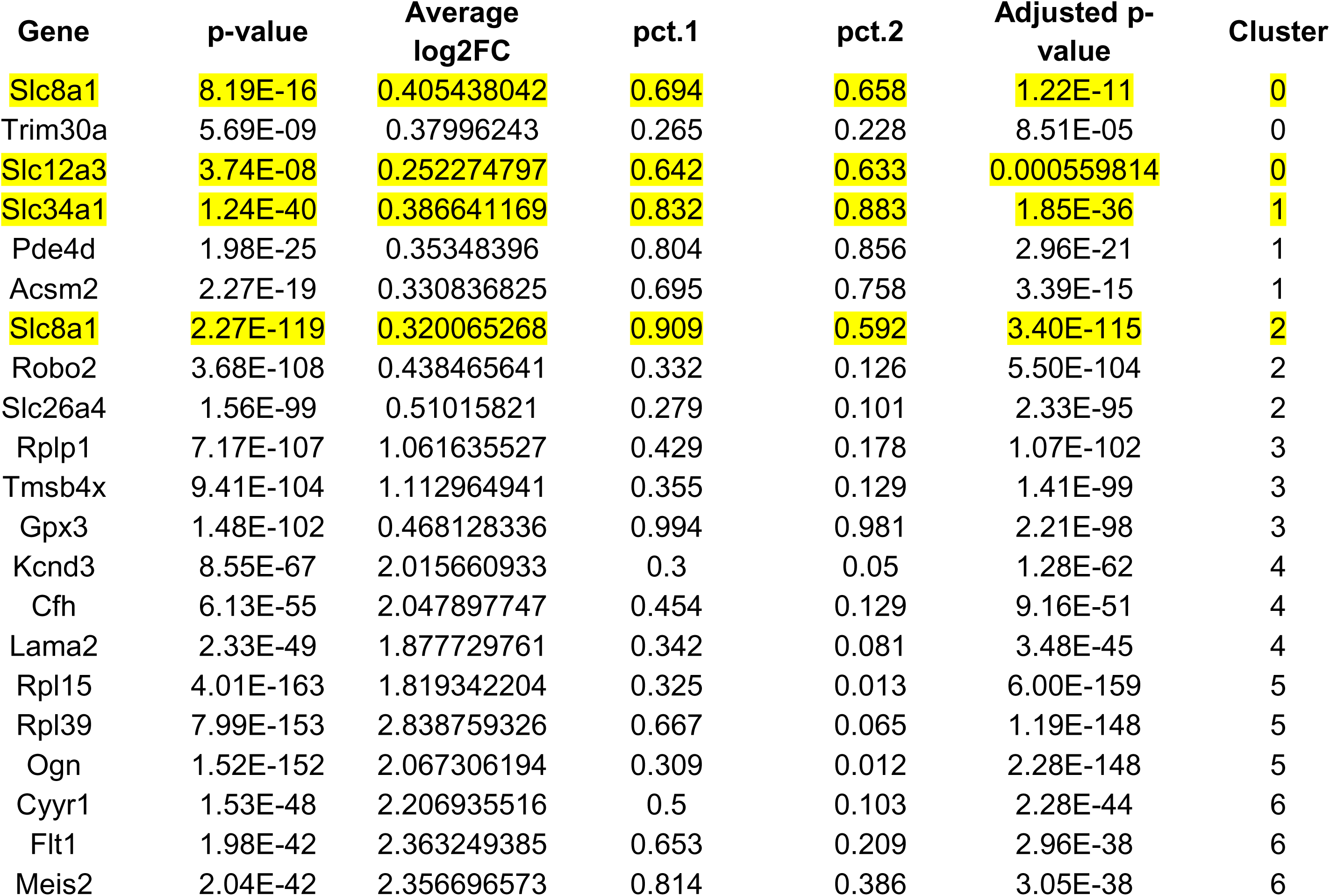
Top 3 cluster marker genes by adjusted p-value for each SC cluster in all samples. Highlighted entries are discussed in the main text.

**Table 2.** Top 3 cluster marker genes by adjusted p-value for each SC cluster in hypertensive samples. Highlighted entries are discussed in the main text. Note: cluster 0 only has two cluster markers.

| Gene | p-value | Average log2FC | pct.1 | pct.2 | Adjusted p-value | Cluster |
| --- | --- | --- | --- | --- | --- | --- |
| S100g | 5.12E-08 | 0.424868363 | 0.151 | 0.267 | 0.00069381 | 0 |
| Magi2 | 7.04E-08 | 0.352053385 | 0.795 | 0.783 | 0.000952946 | 0 |
| **Slc8a1** | **2.46E-33** | **0.354148617** | **0.851** | **0.532** | **3.33E-29** | **1** |
| Mecom | 1.51E-26 | 0.351204224 | 0.753 | 0.43 | 2.04E-22 | 1 |
| Egf | 4.05E-26 | 0.342384186 | 0.514 | 0.274 | 5.48E-22 | 1 |
| Zbtb20 | 6.01E-49 | 0.677214385 | 0.944 | 0.929 | 8.14E-45 | 2 |
| Pde4d | 2.56E-48 | 0.798521338 | 0.89 | 0.867 | 3.47E-44 | 2 |
| Trabd2b | 2.36E-45 | 1.098248484 | 0.755 | 0.674 | 3.19E-41 | 2 |
| Lsamp | 3.28E-50 | 0.428387753 | 0.431 | 0.095 | 4.44E-46 | 3 |
| Pck1 | 1.05E-49 | 0.38912137 | 0.454 | 0.103 | 1.42E-45 | 3 |
| Plce1 | 4.53E-47 | 0.416402462 | 0.385 | 0.082 | 6.13E-43 | 3 |
| D630023F18Rik | 1.59E-38 | 2.36097998 | 0.435 | 0.094 | 2.16E-34 | 4 |
| Atxn1 | 9.58E-34 | 1.585935575 | 0.879 | 0.634 | 1.30E-29 | 4 |
| Slc5a12 | 1.03E-32 | 2.249477306 | 0.629 | 0.249 | 1.40E-28 | 4 |
| Cfh | 5.87E-22 | 2.585559246 | 0.688 | 0.139 | 7.94E-18 | 5 |
| S100a6 | 1.73E-19 | 3.135585438 | 0.469 | 0.07 | 2.35E-15 | 5 |
| Myl9 | 3.93E-17 | 2.483976048 | 0.406 | 0.059 | 5.32E-13 | 5 |

**Table 3.**
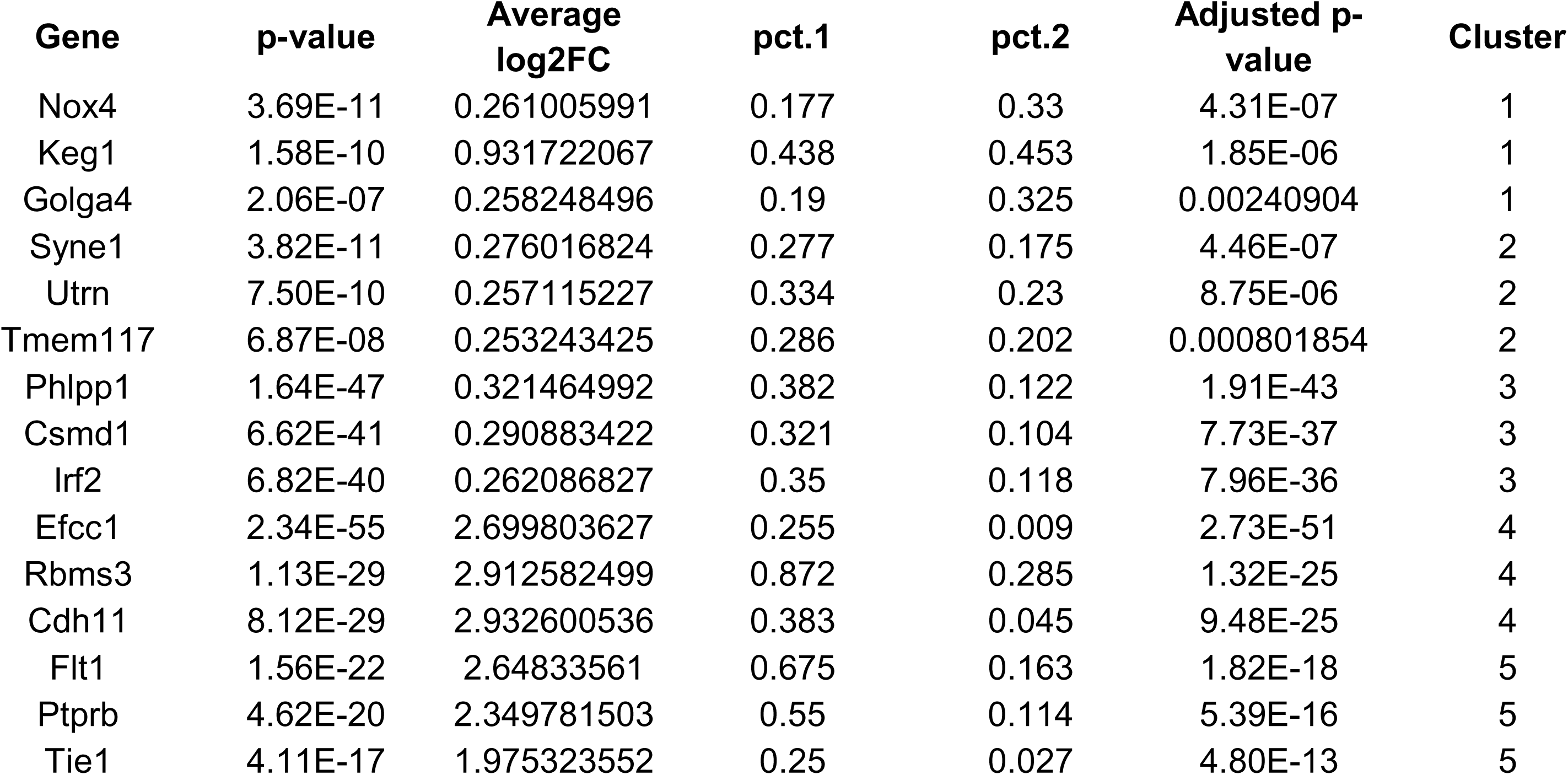
Top 3 cluster marker genes by adjusted p-value for each SC cluster in normotensive samples. Note: cluster 0 has no cluster markers.

### Renal SC-specific gene expression alterations in HTN

To maintain consistency in our approach across the entire data set, we applied the same methods and parameters and again utilized a merged Ctrl group for LECs and SCs and a p-value threshold of <0.01 for significance for all comparisons in this paper. There were a total of 299 differentially expressed genes (DEGs) (254 upregulated, 45 downregulated) when comparing SCs from all HTN samples against the SCs from the Ctrl samples (Table 4 and “DEG” file Table S6). SCs from the A2HTN group had 441 DEGs (383 upregulated, 58 downregulated) compared to Ctrl SCs while SCs from the SSHTN group had 286 (271 upregulated, 15 downregulated) (“DEG” file Tables S7-8). Comparison of the SCs from each HTN group yielded 275 DEGs (46 upregulated in A2HTN, 229 upregulated in SSHTN) (“DEG” file Table S9).

**Table 4.**
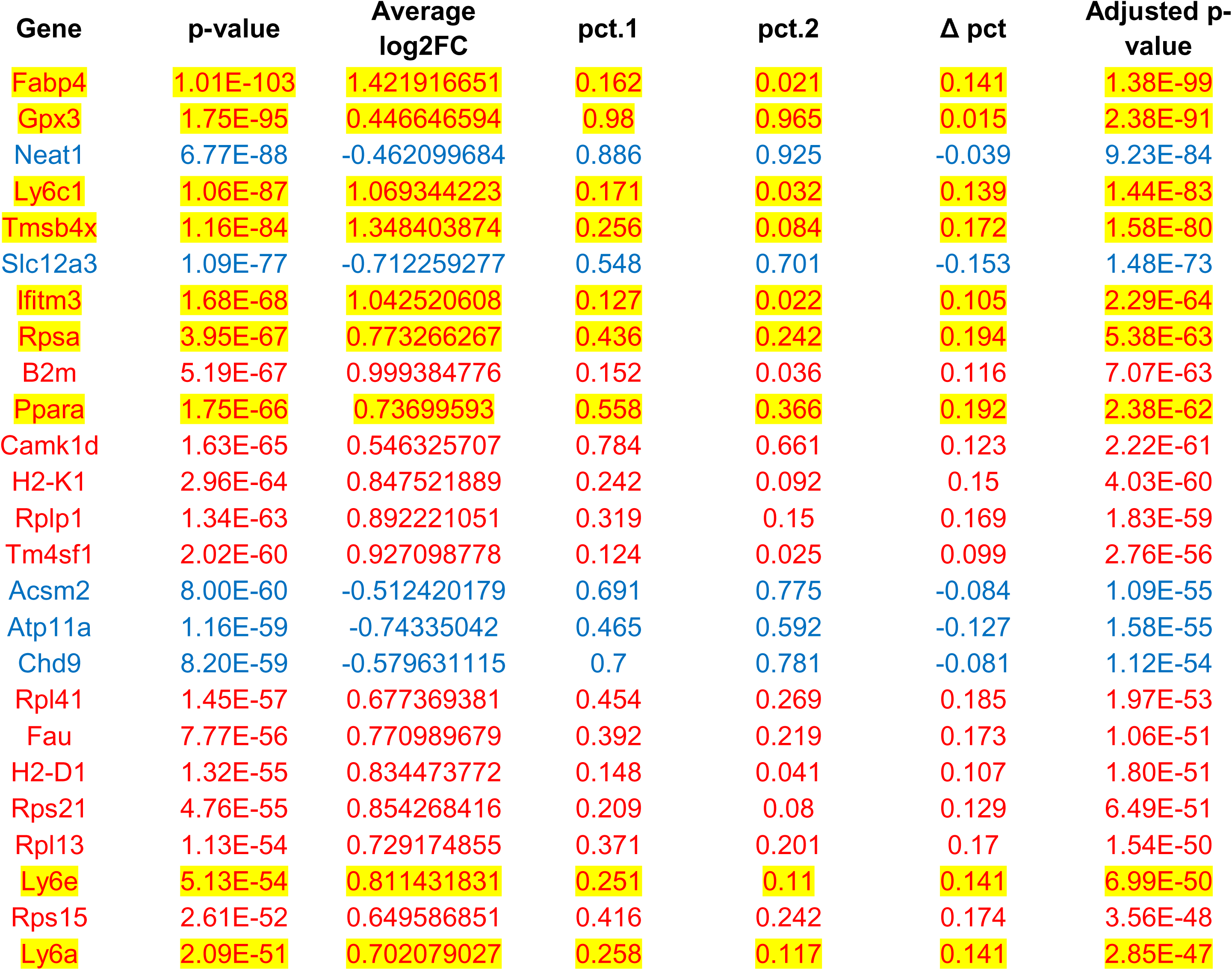
Top 25 differentially expressed genes by adjusted p-value in hypertensive SCs versus normotensive SCs. Highlighted entries are discussed in the main text. (Upregulated genes are shown in red and downregulated genes are shown in blue)

Genes related to cellular growth, proliferation, and survival were among the top upregulated genes in HTN SCs compared to Ctrl SCs, including fatty acid binding protein 4 (*Fabp4*) (168% increase), glutathione peroxidase (*Gpx3*) (36% increase), ribosomal protein SA (*Rpsa*) (71% increase), and peroxisome proliferator activated receptor alpha (*Ppara*) (67% increase).^37–44^ Genes involved in hematopoietic stem cell (HSC) and progenitor stromal cell function were also upregulated, such as *Gpx3*, thymosine beta 4 X-linked (*Tmsb4x*) (155% increase), interferon induced transmembrane protein 3 (*Ifitm3*) (106% increase), and lymphocyte antigen 6 genes (specifically *Ly6a*, *Ly6c1*, and *Ly6e*) as well as multiple ribosomal protein L and S genes.^45–53^

In A2HTN, SCs show a notable increase in MHC class I axis genes, such as *H2-D1* (141% increase), *H2-K1* (118% increase), and beta-2-microglobulin (*B2m*) (156% increase), which are all involved in antigen presentation and inflammatory activation.^54,55^ Matrix gla protein (Mgp) (136% increase) helps to prevent calcification during cardiovascular dysfunction and can suppress the proliferation of T cells and production of inflammatory cytokines.^56^ In SSHTN, SCs have a less pronounced inflammatory response than A2HTN SCs and instead have increases in podoctye (*Magi2* and *Podxl*) and proximal tubule (*Kap*) marker genes, potentially in response to different levels of pressure and damage.

To better understand the connections and distinctions between LECs and SCs in HTN, we compared the two cell types within each group (Tables 5-6 and “DEG” file Tables S10-13). A total of 3636 DEGs (2359 upregulated in LECs, 1277 upregulated in SCs) were identified in the merged HTN group, with similar numbers in A2HTN (3793 [2565 upregulated in LECs, 1228 upregulated in SCs]) and SSHTN (3449 DEGs [2083 upregulated in LECs, 1366 upregulated in SCs]). In contrast, the Ctrl group only had 2194 DEGs (1838 upregulated in LECs, 356 upregulated in SCs). Endothelial genes, including *Sox18*, *Cd34*, and *Ramp2* were consistently and significantly enriched in LECs, while nephron and stem markers, like *Pkhd1*, *Slc34a1*, *Zbtb20*, and *Esrrg*, were reliably upregulated in SCs across groups.^15,30,57^ While sodium-phosphate transport protein 2A (*Slc34a1*) is not significantly altered across any groups in SCs, it is significantly downregulated in LECs in HTN groups, which may be a response to the deleterious effects of excess phosphate and how it can worsen BP.^58,59^ When measured against the dynamic responses that LECs undergo in response to HTN, the SCs demonstrated a relatively muted inflammatory response.^25^

**Table 5.** Top 25 differentially expressed genes by adjusted p-value in hypertensive LECs versus hypertensive SCs. Highlighted entries are discussed in the main text. (Upregulated genes are shown in red and downregulated genes are shown in blue)

| Gene | p-value | Average log2FC | pct.1 | pct.2 | Δ pct | Adjusted p-value |
| --- | --- | --- | --- | --- | --- | --- |
| Sox17 | 0 | 1.967406226 | 0.522 | 0.035 | 0.487 | 0 |
| Rpl7 | 0 | 2.266080443 | 0.85 | 0.109 | 0.741 | 0 |
| Pkhd1 | 0 | -3.309582749 | 0.122 | 0.706 | -0.584 | 0 |
| Rpl31 | 0 | 1.144329097 | 0.739 | 0.179 | 0.56 | 0 |
| Fam117b | 0 | 1.404168614 | 0.696 | 0.138 | 0.558 | 0 |
| Eef1b2 | 0 | 1.584290495 | 0.594 | 0.044 | 0.55 | 0 |
| Klf7 | 0 | 1.540044992 | 0.688 | 0.111 | 0.577 | 0 |
| Rpl37a | 0 | 2.823714533 | 0.972 | 0.233 | 0.739 | 0 |
| Igfbp5 | 0 | 3.071313474 | 0.76 | 0.335 | 0.425 | 0 |
| Arpc2 | 0 | 1.223278796 | 0.659 | 0.129 | 0.53 | 0 |
| Col4a3 | 0 | -2.794577116 | 0.278 | 0.698 | -0.42 | 0 |
| Ptma | 0 | 3.236392853 | 0.976 | 0.192 | 0.784 | 0 |
| Snrpe | 0 | 1.302135382 | 0.433 | 0.02 | 0.413 | 0 |
| Ier5 | 0 | 1.392589241 | 0.454 | 0.019 | 0.435 | 0 |
| Pbx1 | 0 | 1.679074662 | 0.942 | 0.66 | 0.282 | 0 |
| Tagln2 | 0 | 1.565084649 | 0.571 | 0.05 | 0.521 | 0 |
| Mndal | 0 | 1.670939409 | 0.686 | 0.097 | 0.589 | 0 |
| Ifi203 | 0 | 2.187538078 | 0.7 | 0.051 | 0.649 | 0 |
| Itpkb | 0 | 1.365714888 | 0.653 | 0.119 | 0.534 | 0 |
| H3f3a | 0 | 2.744017297 | 0.908 | 0.105 | 0.803 | 0 |
| Esrrg | 0 | -3.73398393 | 0.181 | 0.84 | -0.659 | 0 |
| Cd34 | 0 | 2.007313094 | 0.667 | 0.044 | 0.623 | 0 |
| Edf1 | 0 | 1.420845262 | 0.613 | 0.063 | 0.55 | 0 |
| Notch1 | 0 | 1.347692712 | 0.501 | 0.051 | 0.45 | 0 |
| Egfl7 | 0 | 2.200364762 | 0.896 | 0.185 | 0.711 | 0 |

**Table 6.** Top 25 differentially expressed genes by adjusted p-value in normotensive LECs versus normotensive SCs. Highlighted entries are discussed in the main text. (Upregulated genes are shown in red)

| Gene | p-value | Average log2FC | pct.1 | pct.2 | Δ pct | Adjusted p-value |
| --- | --- | --- | --- | --- | --- | --- |
| B2m | 0 | 3.781757989 | 0.71 | 0.036 | 0.674 | 0 |
| Rps21 | 0 | 3.760025785 | 0.834 | 0.08 | 0.754 | 0 |
| Sox18 | 0 | 2.038037479 | 0.382 | 0.001 | 0.381 | 0 |
| Fabp4 | 0 | 5.174695207 | 0.602 | 0.021 | 0.581 | 0 |
| Tm4sf1 | 0 | 3.509439085 | 0.602 | 0.025 | 0.577 | 0 |
| Rps3a1 | 0 | 2.999908582 | 0.749 | 0.051 | 0.698 | 0 |
| Rps27 | 0 | 3.847298094 | 0.857 | 0.081 | 0.776 | 0 |
| S100a6 | 0 | 3.340269304 | 0.552 | 0.019 | 0.533 | 0 |
| Rpl34 | 0 | 3.242340888 | 0.753 | 0.05 | 0.703 | 0 |
| Gng5 | 0 | 2.748071205 | 0.61 | 0.022 | 0.588 | 0 |
| Id3 | 0 | 3.459695872 | 0.61 | 0.01 | 0.6 | 0 |
| Tomm7 | 0 | 2.221006071 | 0.494 | 0.012 | 0.482 | 0 |
| Hspb1 | 0 | 2.787350799 | 0.483 | 0.011 | 0.472 | 0 |
| Gng11 | 0 | 3.76635646 | 0.629 | 0.016 | 0.613 | 0 |
| Rps5 | 0 | 2.991869208 | 0.749 | 0.046 | 0.703 | 0 |
| Rps16 | 0 | 3.25704105 | 0.795 | 0.064 | 0.731 | 0 |
| Rps3 | 0 | 2.990165829 | 0.753 | 0.052 | 0.701 | 0 |
| Cavin3 | 0 | 2.337902956 | 0.44 | 0.005 | 0.435 | 0 |
| Ifitm2 | 0 | 3.278302389 | 0.656 | 0.019 | 0.637 | 0 |
| Ifitm3 | 0 | 4.028734407 | 0.699 | 0.022 | 0.677 | 0 |
| Tcim | 0 | 2.772802656 | 0.483 | 0.01 | 0.473 | 0 |
| Bst2 | 0 | 2.86092282 | 0.498 | 0.008 | 0.49 | 0 |
| Klf2 | 0 | 2.798787291 | 0.51 | 0.008 | 0.502 | 0 |
| Srgn | 0 | 3.184727844 | 0.587 | 0.019 | 0.568 | 0 |
| Ppia | 0 | 3.873196897 | 0.834 | 0.066 | 0.768 | 0 |

### HTN-related pathway activation in SCs

Gene set enrichment analysis (GSEA) of the DEGs from SCs in HTN and Ctrl groups resulted in 51 enriched pathways (50 upregulated, 1 downregulated) from the curated gene set list (M2) of the Molecular Signatures Database and 86 enriched pathways (77 upregulated, 9 downregulated) from the ontology gene set list (M5) (Figure 2 and “GSEA” file Tables S14-15). Many of the enriched M2 pathways reflect an activated cell state through increased metabolism and translation alongside stem cell and stem-related pathways and the enriched M5 pathways also support this. In A2HTN, the 65 enriched M2 pathways (64 upregulated, 1 downregulated) and 52 enriched M5 pathways (52 upregulated, 0 downregulated) show similar increases, but pathways directly and indirectly related to inflammation response and multipotent properties are further accentuated (Figure S5 A&C and “GSEA” file Tables S16-17). SSHTN contrasts this with its 27 enriched M2 pathways (26 upregulated, 1 downregulated) and 17 enriched M5 pathways (13 upregulated, 4 downregulated) that resemble the merged HTN response, but with significantly less enrichment (either positive or negative) (Figure S5 B&D and “GSEA” file Tables S18-19). When contrasting the two models, the 16 enriched M2 pathways and 1 enriched M5 pathway are all upregulated in A2HTN and relate to metabolism, translation, and inflammation response (“GSEA” file Tables S20-21).

**Figure 2.**
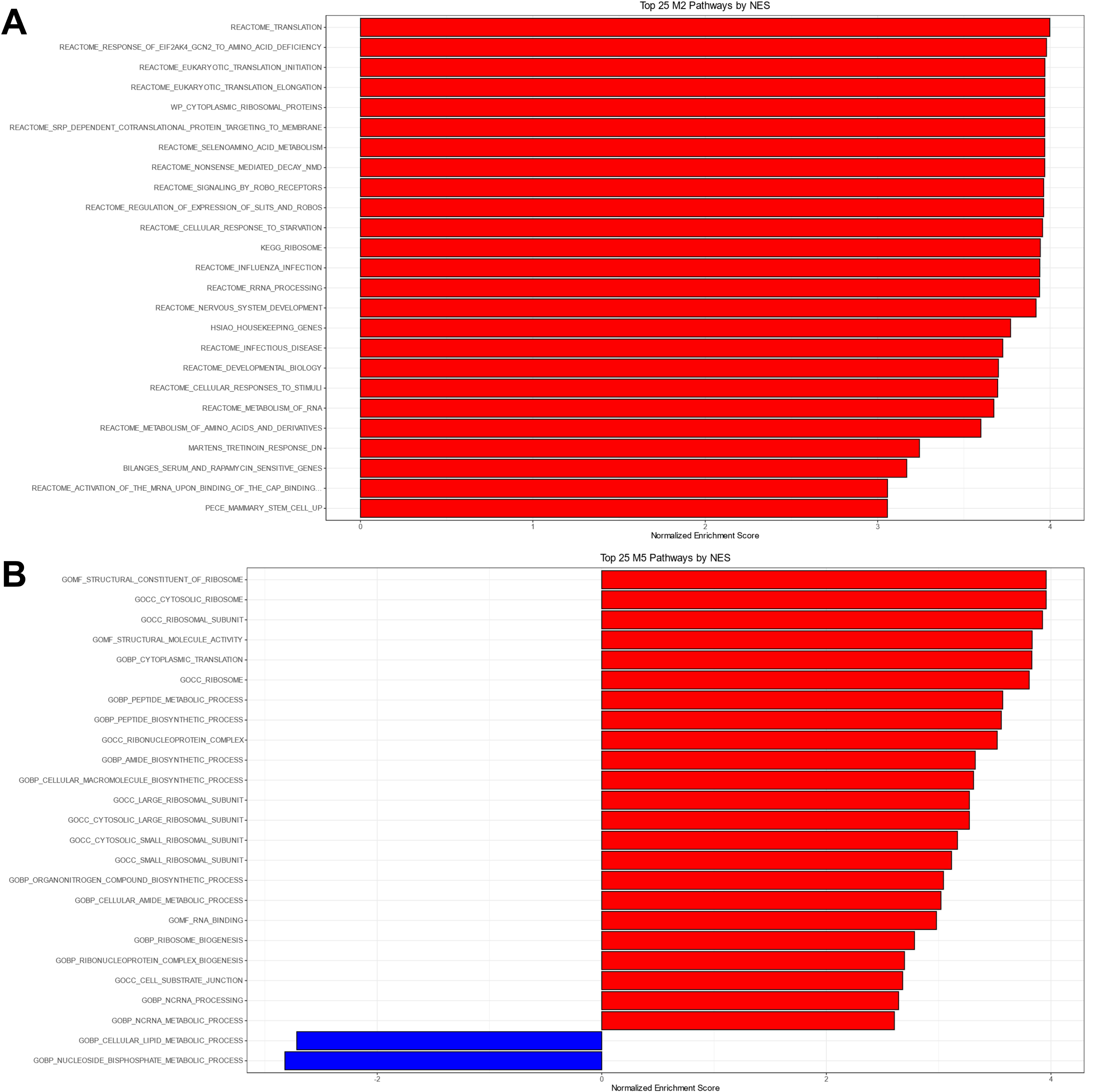
GSEA NES plots for the top 25 M2 (top) and M5 (bottom) pathways by NES from the DEGs between hypertensive SCs and normotensive SCs. The full GSEA results are available in Tables S14-15.

Continuing on with the DEGs from the LEC versus SC comparison in each group, GSEA produced 537 enriched M2 pathways (428 upregulated in LECs, 109 upregulated in SCs) and 415 enriched M5 pathways (247 upregulated in LECs, 168 upregulated in SCs) in the merged HTN group, whereas the Ctrl group had fewer at 507 (425 upregulated in LECs, 82 upregulated in SCs) M2 pathways and 346 (238 upregulated in LECs, 108 upregulated in SCs) M5 pathways (Figure 3 and “GSEA” file Tables S22-25). A2HTN had 646 (509 upregulated in LECs, 137 upregulated in SCs) and 426 (248 upregulated in LECs, 178 upregulated in SCs) enriched M2 and M5 pathways and SSHTN had slightly fewer at 584 (473 upregulated in LECs, 111 upregulated in SCs) and 402 (247 upregulated in LECs, 155 upregulated in SCs) for M2 and M5 pathways (Figure S6 and “GSEA” file Tables S26-29). LECs were upregulated in pathways involving endothelial and leukocyte development, growth, and proliferation, HSCs, inflammation and damage response, and antigen presentation and reactivity. SCs were associated with pathways involving general metabolism, ion transport, kidney-specific cell types, and stem cell markers. In HTN groups, the pattern for each cell type was exaggerated, resulting in additional pathways for each process being upregulated and existing pathways being even further enriched. Slit-Robo signaling, which can enhance lymphangiogenesis, is enriched in LECs over SCs, but it is also enriched in SCs from HTN samples over SCs from controls.^60^ Among the significantly enriched pathways with the lowest NES values in all groups are multiple processes related to vascular endothelial cell identity, potentially indicating some degree of endothelial differentiation in the SCs we identified, even under normotensive conditions.

**Figure 3.**
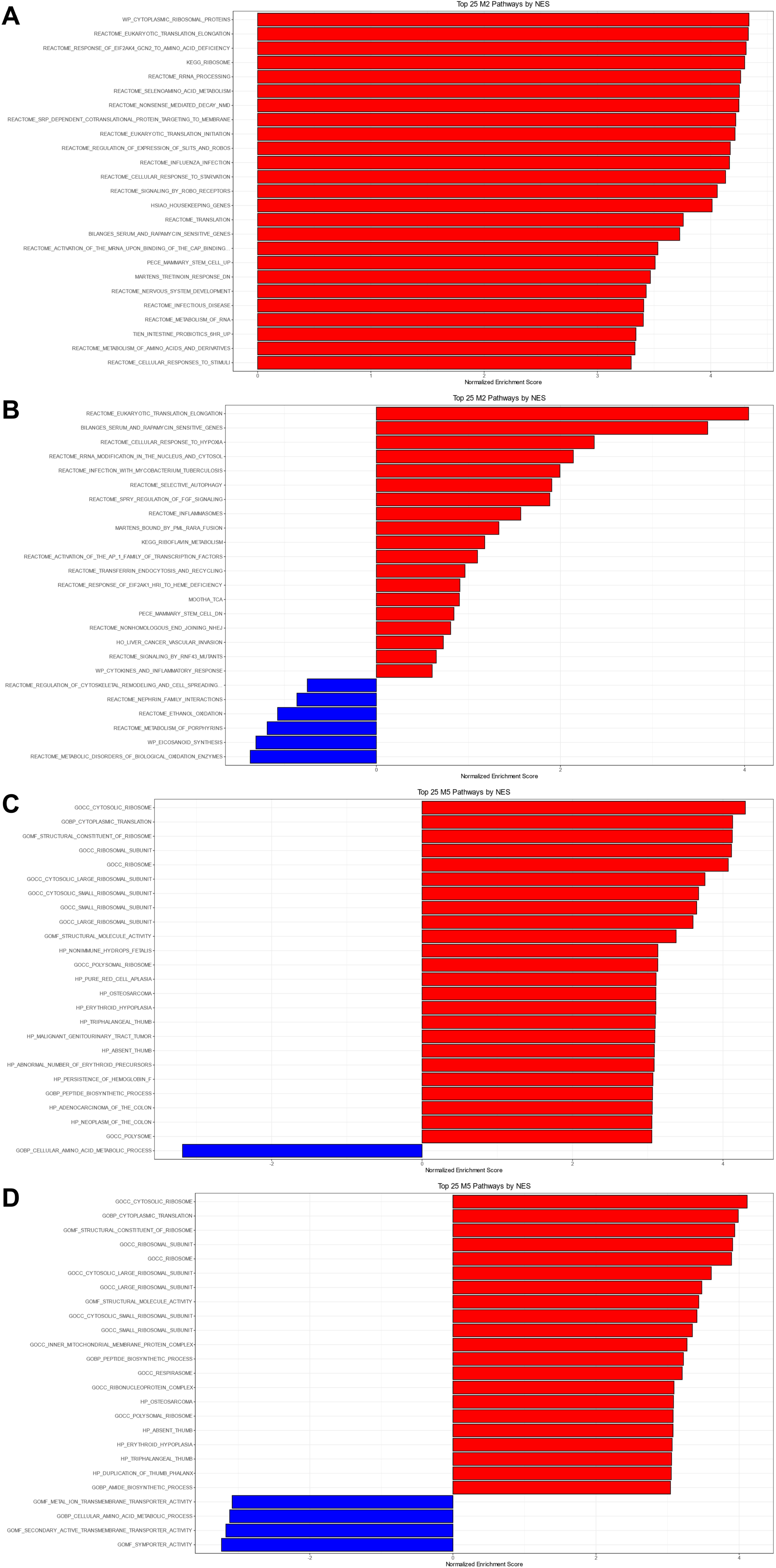
GSEA NES plots for the top 25 M2 (A&B) and M5 (C&D) pathways by NES from the DEGs between hypertensive LECs versus hypertensive SCs (A&C) and normotensive LECs versus normotensive SCs (B&D). Pathways enriched in LECs have positive NES values, while pathways enriched in SCs have negative NES values. The full GSEA results are available in Tables S22-25.

Transcriptional changes promote inflammation and differentiation in SCs due to HTN Transcriptional regulation is vital to defining and maintaining stromal cell identity and we utilized SCENIC to determine which regulons are involved in the changes seen in HTN.^29^ We identified 612 regulons when comparing HTN and Ctrl SCs, with 180 having RSS values >0.5 for HTN SCs, indicating specific or preferential enrichment in the group, and 264 with RSS values >0.5 for Ctrl SCs (“RSS” file Tables S30-33).^61^ While the total number of regulons remained roughly the same in A2HTN and SSHTN versus Ctrl comparisons (608 versus 598 regulons total, respectively), the ratio of HTN to Ctrl regulons shifted in the individual models (117 in HTN and 301 in Ctrl versus 103 in HTN and 308 in Ctrl, respectively) compared to the merged HTN group, with fewer regulons being enriched in HTN groups and more being enriched in the Ctrl group.

Distinctions between LECs and SCs were also evaluated using SCENIC. In the merged HTN group, we found 618 regulons total, with 218 being specific to LECs and 227 that were specific to SCs (“RSS” file Table S34). Each individual HTN model and the Ctrl group all had roughly similar total numbers of regulons, albeit with large differences in number of specific regulons for the two cell types (“RSS” file Tables S35-37). Whereas the merged HTN group had roughly equal numbers, A2HTN leaned more towards SC regulon enrichment and SSHTN showed stronger LEC regulon enrichment. At baseline in the Ctrl group, specific SC regulons outnumber LEC regulons by a wide margin (328 versus 53), which may be reflective of the large number of unique factors that need to remain constitutively active to maintain SC identity and control their proliferation.

Regulons with the largest RSS values per group for each comparison were plotted to show enrichment for each regulon and the distribution of RSS values per group (Figures 4-5 and S7-8) and regulon clustermaps were also generated to showcase the distribution and expression of regulons across samples (Figure S9).

**Figure 4.**
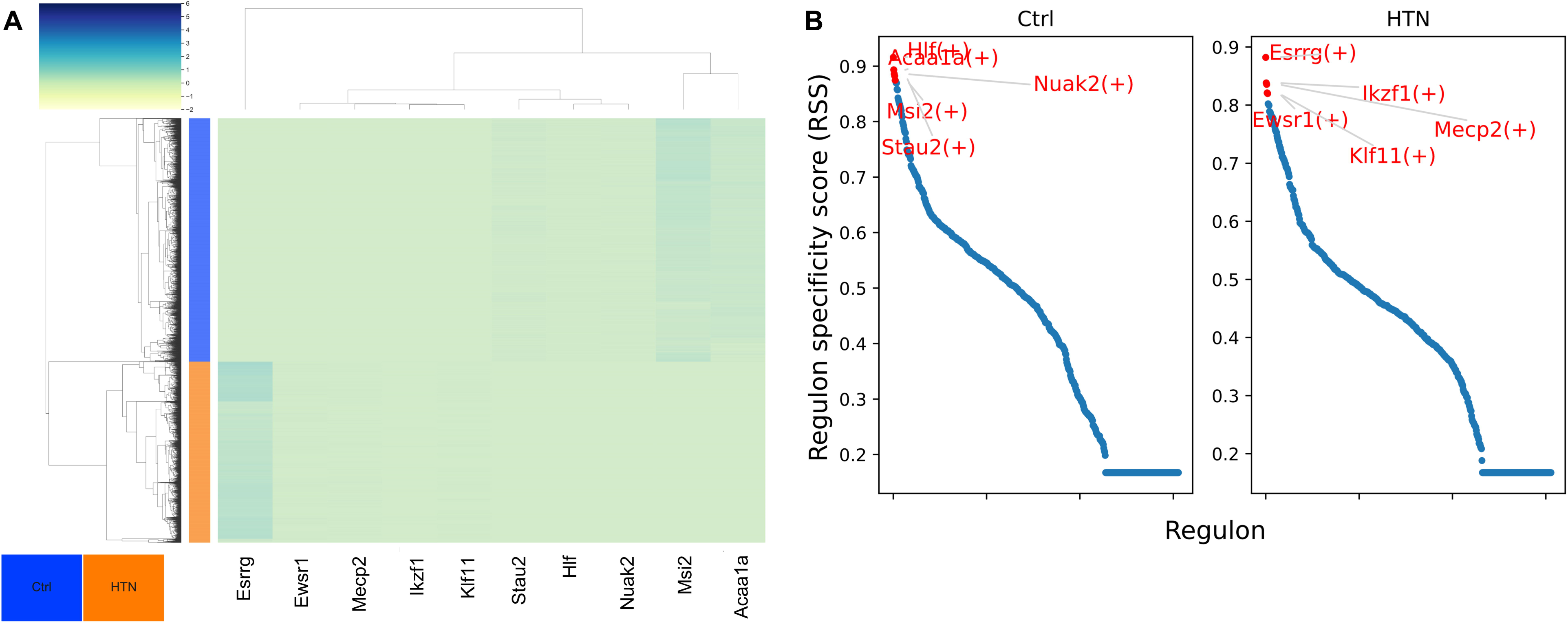
SCENIC RSS heatmap (left) and specificity score plot (right) comparing hypertensive SCs and normotensive SCs. The specificity score plot shows the five regulons with the highest score per group in red and the full RSS list is available in Table S30.

**Figure 5.**
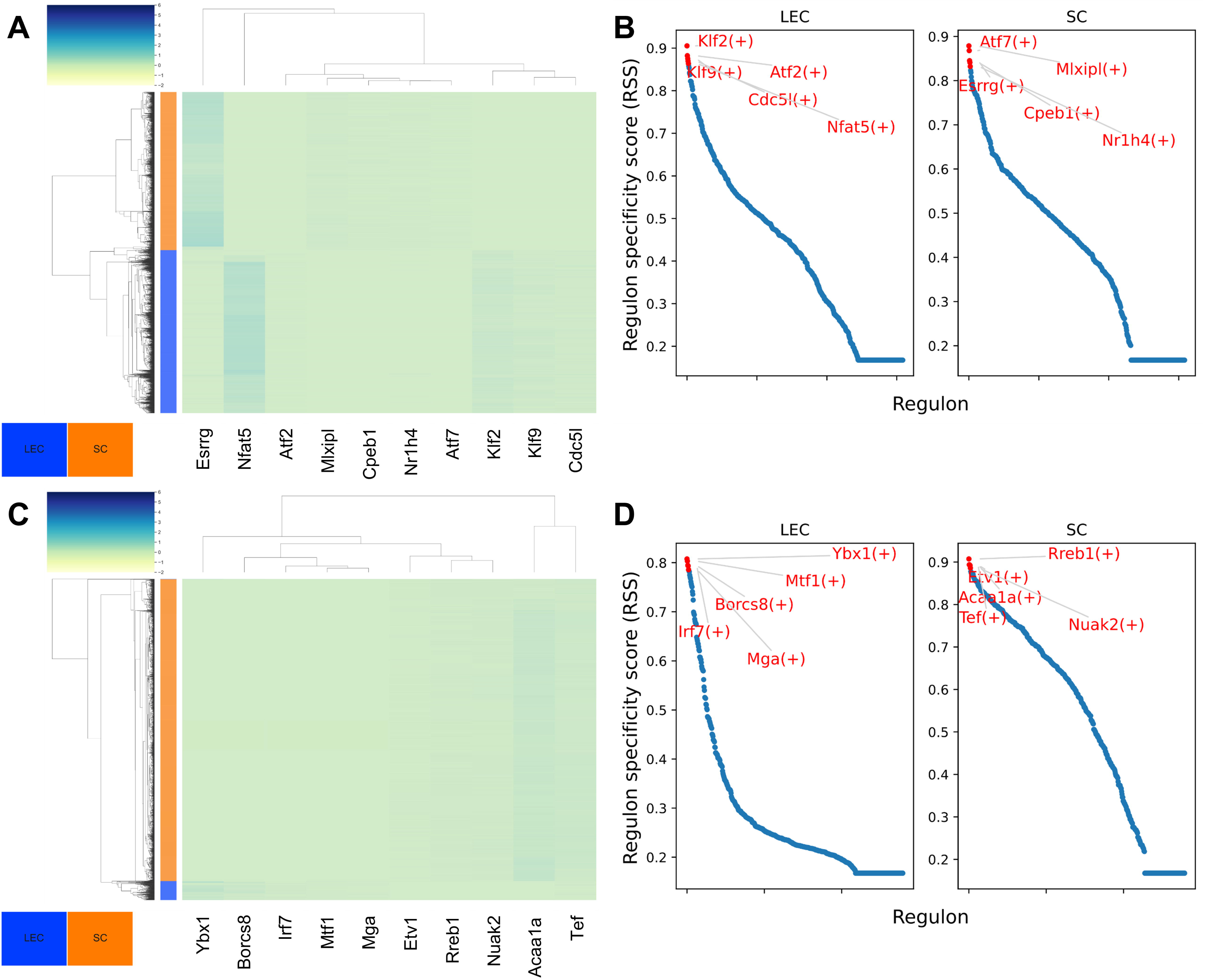
SCENIC RSS heatmap (left) and specificity score plot (right) comparing hypertensive LECs versus hypertensive SCs (top) and normotensive LECs versus normotensive SCs (bottom). The specificity score plot shows the five regulons with the highest score per group in red and the full RSS lists are available in Tables S34-35.

Regulons specific to HTN SCs (versus Ctrl SCs) include *Esrrg*, *Ikzf1*, *Mecp2*, *Ewsr1*, and *Klf11*. *Mecp2* and *Ewsr1* are both involved in stem cell survival through the avoidance or repression of senescence-related pathways and *Mecp2* also positively affect FOXP3, which regulates T-regulatory and T-helper cells.^62–64^ *Esrrg* and *Ikzf1* are markers for differentiation in nephron progenitor cells and HSCs respectively, while *Klf11* has an anti-fibrotic effects in mesenchymal stem cells (MSCs) in the liver and uterus.^30,65–67^ *Hlf*, *Nuak2*, *Msi2*, and *Bptf* are among the regulons that are highly specific to Ctrl SCs, with *Hlf* being associated with undifferentiated multipotent hematopoietic cells and regulation of the early stages of differentiation and *Bptf* maintaining the HSC state through the expression of multiple related transcription factors, such as *Pbx1*, *Meis1*, and *Mn1*.^68,69^ *Nuak2* promotes proliferation, migration, and epithelial-to-mesenchymal transition (EMT) and aids in TGF-β signaling while *Msi2* regulates stem cell maintenance and differentiation and opposes the effects of the differentiating factor *Prox1*.^70–73^

When contrasting HTN LECs and HTN SCs, *Atf2*, *Klf9*, *Cdc5l*, and *Nfat5* were specific to LECs and *Mlxipl*, *Atf7*, *Cpeb1*, *Nr1h4*, and *Mef2a* were specific to SCs. *Cdc5l* is a transcriptional regulator that allows cells to progress through the cell cycle and *Atf2*, *Klf9*, and *Nfat5* are part of the inflammatory response and can promote growth during inflammation.^74–78^ *Atf7* and *Cpeb1* promote and maintain stem-like characteristics and suppress inflammation and proliferation and *Nr1h4* is associated with cilia formation, water reabsorption, and lipid metabolism in the kidney.^79–82^ *Mef2a* promotes survival, growth, and inflammation and can also contribute to muscle regeneration.^83,84^ GWAS studies have implicated the carbohydrate response element binding protein (*Mlxipl*) in liver inflammation, but its role in the kidney, particularly in inflammation, is relatively unknown.^85^

In the Ctrl group, LECs were enriched for *Ybx1*, *Mtf1*, *Irf7*, *Mga*, and *Sox4* while SCs were enriched for *Rreb1*, *Tef, Etv1*, *Creb3l2*, and *Zmiz1*. *Sox4* is expressed by LECs under baseline conditions and *Ybx1* and *Mga* have functions related to cell maintenance and survival.^86–88^ *Irf7* is part of the interferon response family and likely corresponds to the immune-like characteristics of LECs and *Mtf1* can help mitigate damage from hypoxia and oxidative stress.^89,90^ *Etv1* has roles in EMT and stromal growth and survival and *Rreb1* pushes cells towards EMT and maintains stem characteristics, as well as opposing *Prox1*-fueled differentiation (similar to *Msi2*).^91–93^ *Tef* and *Zmiz1* both regulate HSCs to control proliferation and maintain their stem state, whereas *Creb3l2* is associated with podocytes, which may be a result of having a stem-like state in the kidney.^94–96^

## Discussion

### Known and Potential Changes in the Identity of SCs in HTN

To date, most studies involving pluripotent or multipotent cells in HTN have focused on the role of stem cells as a potential therapeutic measure, where they can be isolated, cultured, and injected to provide a net positive effect through BP reduction and cellular regeneration on various forms of HTN.^10–13,15^ Conversely, very little is known regarding which exact mechanisms that these cells, whether they are injected or native to the tissue, use to bring about these effects, with the most common explanations relating to secreted factors, their regenerative potential through proliferation, or their ability to reduce inflammation locally.^97–99^ Here we sought to gain better mechanistic insight into the role of an unusual multipotent renal cell type in HTN through differential gene expression and pathway analysis, as well as the activity of the regulons governing these changes.

The effects of HTN on the kidney can be simply described as damage and inflammation, which are both relevant to well-established roles of cells similar to SCs. Here we have identified transcriptional changes that are reflective of differentiation into multiple distinct cell types, antigen presentation, anti-inflammatory factors, and continued maintenance of a stem-like state. Across both models of HTN, we observed upregulation in growth-related genes and in many of the same genes that are highly expressed in LECs in our samples, as well as an increase in podocyte and tubular epithelium markers, particularly in the SSHTN model. The A2HTN model led to the induction of MHC class I antigen-presenting genes and *Mgp*, indicating that it had a stronger immunological effect on SCs than the SSHTN model, similar to what was seen in LECs. In HTN, SCs had roughly half as many significant DEGs compared to LECs, which may correspond to a weaker overall response to damage and inflammation in the SCs that maintained their identity during HTN.

It is well-known that multipotent cells can replicate and transdifferentiate under stress and we have previously established that there is an increase in CD31+/podoplanin+ LECs in HTN.^6,19,20,100^ Considering how SCs upregulated genes that are associated with LEC clusters in our samples in response to HTN, it is possible that one mechanism behind this increase is the conversion of SCs or their progeny into LECs or LEC-like cells to supplement existing LEC populations in the kidneys during inflammation. This is further supported by the pathways with the lowest enrichment scores when comparing the two cell types being directly or indirectly involved with endothelial growth and development, even in the control samples.

Additionally, the number of markers associated with podocytes and tubular epithelium in SCs from each sample and the overall increase in these markers in HTN, particularly in SSHTN, suggest that these SCs may be associated with these cell types or even capable of differentiating into them, though it is unclear why this would be more prevalent in SSHTN than in A2HTN. This association may be related to glomerular and tubular regeneration after extensive cortical damage in the kidneys, but this damage is known to worsen with increasing BP, so it could be expected to have a greater impact in A2HTN than in SSHTN, though this is not the case.

### SCs Maintain Relative Quiescence at Baseline and in HTN

Despite the serious increase in inflammatory factors and cells present in the kidneys in HTN, the SCs described here seem to be relatively inactive and quiescent when compared to LECs. The majority of the pathway alterations seen in LECs in HTN had negative NES values, meaning that they were significantly downregulated in response to HTN, with most of the upregulated pathways being tied to inflammation and immune-related activity. In SCs, we see the opposite; there are fewer significantly enriched pathways compared to LECs, but nearly all of them are upregulated. Many of the genes, pathways, and regulons we have reported here in SCs from the control samples have a suppressive effect on stem or progenitor cells in terms of their function, growth, and proliferation and these cell types are known to remain relatively inactive throughout adulthood in order to maintain homeostasis and prevent cancerous activity.^97^ This suppressive effect seems to be decreased or shifted to different genes and pathways in HTN to maintain similar overall effects in an altered environment, with many of these HTN-induced genes pushing towards cell survival, differentiation, and supporting regenerative healing.

### Transcriptional Differences Highlight Functional Distinctions between LECs and SCs

When contrasting LEC and SC gene expression across samples, SCs consistently expressed stem and nephron-relevant markers whereas LECs expressed primarily endothelial and leukocyte markers with some genes associated with hematopoietic stem cells. In HTN, LECs demonstrated an immense inflammatory response, with most upregulated genes, pathways, and regulons reflecting inflammation or growth in some way, while SCs showed significantly less enrichment in these same pathways and instead had comparative upregulation in genes involved in metabolism, ion transport and balance, suppression of inflammation and growth, and stem cell survival. Even in control samples, SCs were enriched for genes related to EMT and maintenance of stem cell identity, suppression of proliferation, and kidney-specific cell types. We also identified a number of potential markers for differentiating between LECs and SCs, including *Ifitm2*, *Ifitm3* and *B2m*, which are primarily expressed in LECs in our samples despite *Ifitm3* and *B2m* also being upregulated in SCs in HTN.

## Conclusion

Pluripotent and multipotent cells are known to have far-reaching effects in response to damage and inflammation and can aid in reducing inflammatory activity, improving regeneration, and restoring functionality in tissue. Following up from our previous paper, where we used scRNAseq to analyze changes in CD31+/podoplanin+ cells isolated from the kidneys of hypertensive and normotensive mice generated using the A2HTN and SSHTN models of HTN, we wanted to determine the role and effects of the SC populations that we identified in each sample and utilized the same methods to determine how these cells responded to HTN. There were increases in genes related to growth, inflammation, podocytes, tubular epithelium, and in genes that were highly expressed in the LEC populations described in the companion paper, indicating a close association with or differentiation into these cell types. We also noted that many of the genes expressed by SCs in our control samples were related to the maintenance of stem cell identity and suppression of growth and that these genes are replaced by others that push toward stem cell proliferation and differentiation in HTN. Functional distinctions between LECs and SCs in each sample were also characterized and help to highlight the divergent pathways that each cell type uses to mitigate the effects of HTN.

Going forward, we intend to study how this SC population impacts renal lymphangiogenesis in HTN to determine if it contributes to the increases that we have described previously, either through differentiation or the addition of various pro-lymphangiogenic factors. Due to the relatively small sample size, more work will need to be done to verify the findings presented here with a larger sample and potentially with different additional markers for cell selection to improve the process of isolating SCs.

We also plan to characterize how injections of adipose and bone-derived MSCs primed with inflammatory cytokines interact with renal LECs and SCs in normotensive and hypertensive mice to determine if they can be utilized therapeutically to reduce BP and improve kidney regeneration and function. Further insight into how SCs impact the kidneys during the pathogenesis of HTN could help to create new ways to treat serious cases of HTN and these findings may provide additional understanding into poorly understood mechanisms of renal repair, including potential markers for differing levels of damage and subsequent repair in the kidney as a result of pressure-related injury.

---

Figure S1 – Comparison of cell types identified in our samples versus control/sham mouse kidney samples from NCBI GEO produced by Janosevic et al., Conway et al., and Miao et al. A – UMAP plot showing cells by cell type or by source. B – UMAP plot showing samples after reclustering. C – Feature plots of samples using cell type specific genes derived from previous kidney scRNAseq studies. Included genes are markers for podocytes (*Nphs1*, *Nphs2*), proximal tubule cells (*Slc27a2*, *Slc34a1*, *Lrp2*, *Slc5a2*, *Slc22a30*), distal tubule cells (*Slc12a3*, *Umod*), Loop of Henle cells (*Umod*, *Slc8a1*, *Slc12a1*, *Aqp2*, *Slc26a4*), parietal epithelial and juxtaglomerular cells (*Pax8*, *Ren1*), mesangial cells (*Plvap*, *Prkca*, *Nt5e*, *Pdgfrb*), endothelial cells (*Pecam1*, *Flt1*, *Kdr*, *Nrp1*, *Egfl7*), as well as the activation marker *Cd38*.

Figure S2 – Feature plot of SCs from all samples showcasing expression of cell type specific genes derived from previous kidney scRNAseq studies. Included genes are markers for podocytes (*Nphs1*, *Nphs2*), proximal tubule cells (*Slc27a2*, *Slc34a1*, *Lrp2*, *Slc5a2*, *Slc22a30*), distal tubule cells (*Slc12a3*, *Umod*), Loop of Henle cells (*Umod*, *Slc8a1*, *Slc12a1*, *Aqp2*, *Slc26a4*), parietal epithelial and juxtaglomerular cells (*Pax8*, *Ren1*), mesangial cells (*Plvap*, *Prkca*, *Nt5e*, *Pdgfrb*), and endothelial cells (*Pecam1*, *Flt1*, *Kdr*, *Nrp1*, *Egfl7*).

Figure S3 – TSNE cluster plot and feature plot of SCs from hypertensive (top row) and normotensive (bottom row) samples.

Figure S4 – TSNE cluster plot and feature plot of SCs from A2HTN (top row) and SSHTN (bottom row) samples.

Figure S5 – GSEA NES plots for the top 25 M2 (top) and M5 (bottom) pathways by NES from the DEGs between A2HTN SCs and normotensive SCs (left) and SSHTN SCs and normotensive SCs (right). The full GSEA results are available in Tables S16-19.

Figure S6 – GSEA NES plots for the top 25 M2 (top) and M5 (bottom) pathways by NES from the DEGs between A2HTN LECs versus A2HTN SCs (left) and SSHTN LECs versus SSHTN SCs (right). Pathways enriched in LECs have positive NES values, while pathways enriched in SCs have negative NES values. The full GSEA results are available in Tables S26-29.

Figure S7 – SCENIC RSS heatmap (left) and specificity score plot (right) comparing A2HTN SCs and normotensive SCs (top) and SSHTN SCs and normotensive SCs (bottom). The specificity score plot shows the five regulons with the highest score per group in red and the full RSS lists are available in Table S31-32.

Figure S8 – SCENIC RSS heatmap (left) and specificity score plot (right) comparing A2HTN LECs versus A2HTN SCs (top) and SSHTN LECs versus SSHTN SCs (bottom). The specificity score plot shows the five regulons with the highest score per group in red and the full RSS lists are available in Tables S36-37.

Figure S9 – SCENIC regulon clustermaps showing regulon enrichment throughout all SCs in an entire sample. A – all samples, B – hypertensive samples, C – control samples, D – A2HTN sample, E – SSHTN sample.

## Declaration

Funding: This work was funded by NIH R01 (DK120493) to B.M. Mitchell and Texas A&M University T3 to J.M. Rutkowski.

## Acknowledgement

N/A

## Author Contributions

Justin G. McDermott-conceptualization, data curation, formal analysis, investigation, methodology, project administration, resources, software, writing-original draft, writing-review and editing; Bethany L. Goodlett-data curation, investigation; Shobana Navaneethabalakrishnan-data curation, investigation; Joseph M. Rutkowski-conceptualization, funding acquisition, project administration; Brett M. Mitchell-conceptualization, funding acquisition, project administration, resources, writing-review and editing

## Conflict of Interest

The authors have nothing to disclose. Consent: N/A

Ethics: IACUC Animal Use Protocol TAMU #2022-0083

Data Availability: Raw and processed sequencing data are available through NCBI GEO (GSE236410). All scripts for analysis and their direct results are available upon reasonable request to the corresponding author.

## Notes

https://www.ncbi.nlm.nih.gov/geo/query/acc.cgi?acc=GSE236410

### Competing Interest Statement

The authors have declared no competing interest.

